# Discovery of N-Glycan Outer Chain like α-(1→6)-linked Mannan Structures in *Aspergillus fumigatus* Mycelium

**DOI:** 10.1101/2023.12.08.570825

**Authors:** Chihiro Kadooka, Yutaka Tanaka, Rintaro Kishida, Daisuke Hira, Takuji Oka

**Author notes:** Address correspondence to Takuji Oka.

## Abstract

The cellular surface of the pathogenic filamentous fungus *Aspergillus fumigatus* is enveloped in a mannose layer, featuring well-established fungal-type galactomannan and O-mannose type galactomannan. This study reports the discovery of cell wall component in the mycelium that resemble N-glycan outer chains found in yeast. The glycosyltransferases involved in their biosynthesis in *A. fumigatus* were identified, with a focus on two key α-(1→2)-mannosyltransferases, Mnn2 and Mnn5, and two crucial α-(1→6)-mannosyltransferases, Mnn9 and Van1. *In vitro* examination revealed the roles of Mnn2 and Mnn5 in transferring α-(1→2)-mannosyl residues. ^1^H-NMR analysis of cell wall extracts from the Δ*mnn2*Δ*mnn5* strain indicated the existence of an α-(1→6)-linked mannan backbone in the mycelium, with Mnn2 and Mnn5 adding α-(1→2)-mannosyl residues to this backbone. The α-(1→6)-linked mannan-derived chemical shift was absent in strains where *mnn9* or *van1* was disrupted in the parental Δ*mnn2*Δ*mnn5* strain. Co-expressed Mnn9 and Van1 functioned as α-(1→6)-linked mannan polymerases in heterodimers, suggesting their crucial role in biosynthesizing the α-(1→6)-linked mannan backbone. Disruptions in these mannosyltransferases did not affect fungal-type galactomannan biosynthesis. The study provides insights into the complexity of fungal cell wall architecture and a comprehensive understanding of mannan biosynthesis in *A. fumigatus,* including the presence of outer chain like mannan structures and their potential role in cell wall integrity.

**Importance:** This study unravels the complexities of mannan biosynthesis in *A. fumigatus,* a key area for antifungal drug discovery. It reveals the presence of outer chain like mannan structures, offering fresh insights into the fungal cell wall’s design. Key enzymes Mnn2, Mnn5, Mnn9, and Van1 are instrumental in this process, with Mnn2 and Mnn5 adding specific mannose residues and Mnn9 and Van1 assembling the outer chain like structures. While fungal-type galactomannan’spresence in the cell wall is known, the existence of outer chain like mannan adds a new dimension to our understanding. This intricate web of mannan biosynthesis opens avenues for further exploration and enhances our understanding of fungal cell wall dynamics, paving the way for targeted drug development.

## Introduction

Mannan plays a crucial role in maintaining the cell wall structure in yeast and filamentous fungi. In *Aspergillus fumigatus*, a prominent pathogenic fungus causing invasive pulmonary aspergillosis, mannose-containing sugar chains are integral components found in N-and O-linked glycans on various proteins, glycosylphosphatidylinositol (GPI) anchor, mannosylinositol phosphorylceramides (MIPC), and fungal-type galactomannan (FTGM) (Fontaine and Latgé. 2020, Latgé. 2009. Kudoh et al. 2015, Oka. 2018, Latgé. 2023). Studies from our research group and others have revealed that disrupting *cmsA*/*ktr4*, an α-(1→2)-mannosyltransferase encoded-encoding gene responsible for FTGM α-core mannan synthesis, leads to abnormal colony morphology and a reduction in infectious capacity in *A. fumigatus* (Onoue et al. 2018, Henry et al. 2019). Furthermore, the double disruption of *pmt4* and *pmt1*, encoding protein O-mannosyltransferases,is synthetic lethal, emphasizing the essential role of protein O-mannosylation in hyphal growth through the maintenance of cell surface and secreted proteins (Mouyna et al. 2010). Additionally, disrupting *mnt1*, an α-(1→2)-mannosyltransferase gene responsible for decorating second mannosyl residues to protein O-mannose type galactomannan (OMGM) and elongating high mannose type N-glycan core chain, results in a pale cell wall and hypovirulence (Wagener et al. 2008, Kadooka et al. 2022a). Therefore, comprehensively understanding mannan biosynthesis in *A. fumigatus* is crucial for advancing novel antifungal drug discovery.

In *Saccharomyces cerevisiae*, N-glycans attached to proteins exhibit extensive mannan structures, referred to as outer chains, consisting of up to 200 mannosyl residues (Herscovics and Orlean. 1993). The biosynthesis of outer chain structures is well-studied in *S. cerevisiae*. The initial reaction is catalyzed by the specific α-(1→6)-mannosyltransferase *Sc*Och1p (Nakayama et al. 1992). Subsequently, the elongation of at least 10 mannosyl residues is facilitated by Mannan polymerase I (M-Pol I), a heterodimeric complex of two α-(1→6)-mannosyltransferases, *Sc*Mnn9p and *Sc*Van1p (Ballou et al. 1980, Hashimoto and Yoda. 1997, Jungmann and Munro. 1998). The α-(1→6)-linked mannan backbone is further extended by another mannan polymerase complex, M-Pol II, consisting of α-(1→6)-mannosyltransferases *Sc*Mnn9p, *Sc*Anp1p, *Sc*Mnn10p, *Sc*Mnn11p, and *Sc*Hoc1p (Jungmann et al. 1999, Kojima et al. 1999). Additionally, α-(1→2)-, α-(1→3)-, and mannose-6 phosphate side chains are decorated by α-(1→2)-mannosyltransferases *Sc*Mnn2p and *Sc*Mnn5p (Rayner and Munro. 1998), α-(1→3)-mannosyltransferase Mnn1p (Wiggins and Munro. 1998), and mannose-6 phosphate transferase *Sc*Mnn4p and *Sc*Mnn14p (Odani et al. 1997, Kim et al. 2017). The visualization of the outer chain α-(1→6)-linked mannan core backbone in a *Scmnn2* mutant strain, for the first time, identified *Sc*Mnn2p as an α-(1→2)-mannosyltransferase responsible for the initial elongation of α-(1→2)-mannosyl residues in the side chain of N-glycan outer chain (Ballou. 1975). Subsequently, *Sc*Mnn5p decorates second α-(1→2)-mannosyl residues to the side chain. Finally, the N-glycan outer chain structures are fully biosynthesized in *S. cerevisiae* through the decoration of α-(1→3)-mannosyl residues and mannose 6-phosphate onto (1→2)-mannosyl residues, dependent on *Sc*Mnn1p, *Sc*Mnn14p, and *Sc*Mnn4p.

The initial specific α-(1→6)-mannosyltransferase, *Sc*Och1p serves as a pivotal enzyme in the biosynthesis of outer chain structures. Analyses of *Sc*Och1p orthologues in various model and pathogenic fungi have been conducted. In the pathogenic yeast *Candida albicans*, disruption of *Caoch1* has been reported to strongly induce growth defects and reduce infection survival rates (Bates et al. 2005). Similarly, the *och-1* disruptant in exhibited severe growth defects and abnormal conidial formations, suggesting the presence of N-glycan outer chain structures not only in yeast but also in filamentous fungi (Maddi and Free. 2010). Contrastingly, in *A. fumigatus*, disruption of the *och1* gene resulted in only growth defects and abnormal conidia formations under calcium stress conditions (Kotz et al. 2010). Another research group reported that a quadruple disruptant of four paralogous och1 genes (*och1-1∼1-4*) did not exhibit any phenotypes in *A. fumigatus*, suggesting a potential divergence in mannan biosynthesis pathways between filamentous fungi and yeast (Lambou et al. 2010). In a study by Henry et al. (2016), the Δ*11* strain (*mnn9*, *van1*, *anp1*, *mnn10*, *mnn11*, *mnn2*, *mnn5*, and *och1* orthologue genes multiple-disruptant) showed a growth defect and reduced cell wall thickness and alkali-soluble mannan content, indicating the possible roles of these mannosyltransferases in biosynthesizing specific mannan structures. Du et al. (2019) reported the requirement of the Mnn9p orthologue for cell wall integrity through cell wall mannan and/or mannoprotein biosynthesis in *A. fumigatus*. Additionally, a recent study by Liu et al. (2023) revealed an unknown glycan structure named G3Man in *A. fumigatus* conidia, specifically present in the conidia, implicating its biosynthesis in the GT62 family genes triple disruptant (including *mnn9*, *van1,* and *anpA*) and GT32 family genes quadruple disruptant (including och1-4). Unlike the yeast outer chain, G3Man lacks α-(1→2)-mannosyl side chains on the α-(1→6)-linked mannan backbone but features side chains of galactose, glucose, and N-acetylglucosamine. However, the functions of Mnn2 and Mnn5 homologs in filamentous fungi and the presence and importance of the outer chain in mycelia remain unclear.

In our study, we specifically focused on two putative α-(1→2)-mannosyltransferase, Mnn2 and Mnn5. Mannosyltransferase and substrate-specific mannosidase assays, utilizing recombinant Mnn2 and Mnn5, revealed their shared α-(1→2)-mannosyltransferase activity *in vitro*. While single gene disruptants of mnn2 or mnn5 showed no discernible phenotype, the mnn2/mnn5 double disruptant exhibited a growth defect and abnormal conidial formations. ^1^H-NMR analysis of mannoproteins from the mnn2/mnn5 double disruptant provided compelling evidence for the presence of an outer chain like α-(1→6)-linked mannan backbone in *A. fumigatus* mycelia. Furthermore, Mnn9 and Van1 were identified as key enzymes in the biosynthesis of the α-(1→6)-linked mannan backbone. In our *in vitro* experiments, recombinant Mnn9 and Van1 were shown to form a heterodimer and catalyze the synthesis of α-(1→6)-mannose polymers.

## Results

### Mannosyltransferase activities of Mnn2 and Mnn5 *in vitro*

In our investigation of the mannosyltransferase activities of Mnn2 (AFUB_093840) and Mnn5 (AFUB_060800) from *A. fumigatus*, we produced individual N-terminal 6×Histagged recombinant proteins using a bacterial expression system. Mnn2 and Mnn5 were expressed without the presumed transmembrane domains, spanning amino acid residues 1-30 and 1-29, respectively. Analysis of the purified recombinant proteins through sodium dodecyl sulfate-polyacrylamide gel electrophoresis (SDS-PAGE) revealed bands close to their predicted molecular weights of 53 and 55 kDa, respectively (Fig. 1A). The yields of the recombinant proteins were 0.88 mg/L for Mnn2 and 1.25 mg/L for Mnn5. Subsequently, we assessed mannosyltransferase activity at 30°C for 16 h using recombinant Mnn2 and Mnn5 (0.2 µg/µL each). The reaction involved *p*-nitrophenyl α-D-mannopyranoside (α-Man-pNP, 1.5 mM) as the acceptor substrate, guanosine diphosphate-α-D-mannose (GDP-Man, 5 mM) as the donor substrate, and 0.5 mM Mn2+ as the metal cofactor. Fractions with heat-inactivated Mnn2 and Mnn5 exhibited no new peaks (Fig. 1B, upper panels). Conversely, the fractions containing Mnn2 or Mnn5 revealed new products (referred to as Mnn2-product and Mnn5-product) at 18.0 min (Fig. 1B, middle and bottom panels). To elucidate the chemical structures of Mnn2-product and Mnn5-product, we isolated each peak and digested them using a substrate-specific mannosidase. Recombinant MsdS/MsdC, an α-(1→2)-specific mannosidase from *A. fumigatus*, was employed to convert Mnn2-product and Mnn5-product to α-Man-pNP (Fig. 1C). These findings provide compelling evidence supporting the role of Mnn2 and Mnn5 as α-(1→2)-mannosyltransferases.

**Fig. 1.**
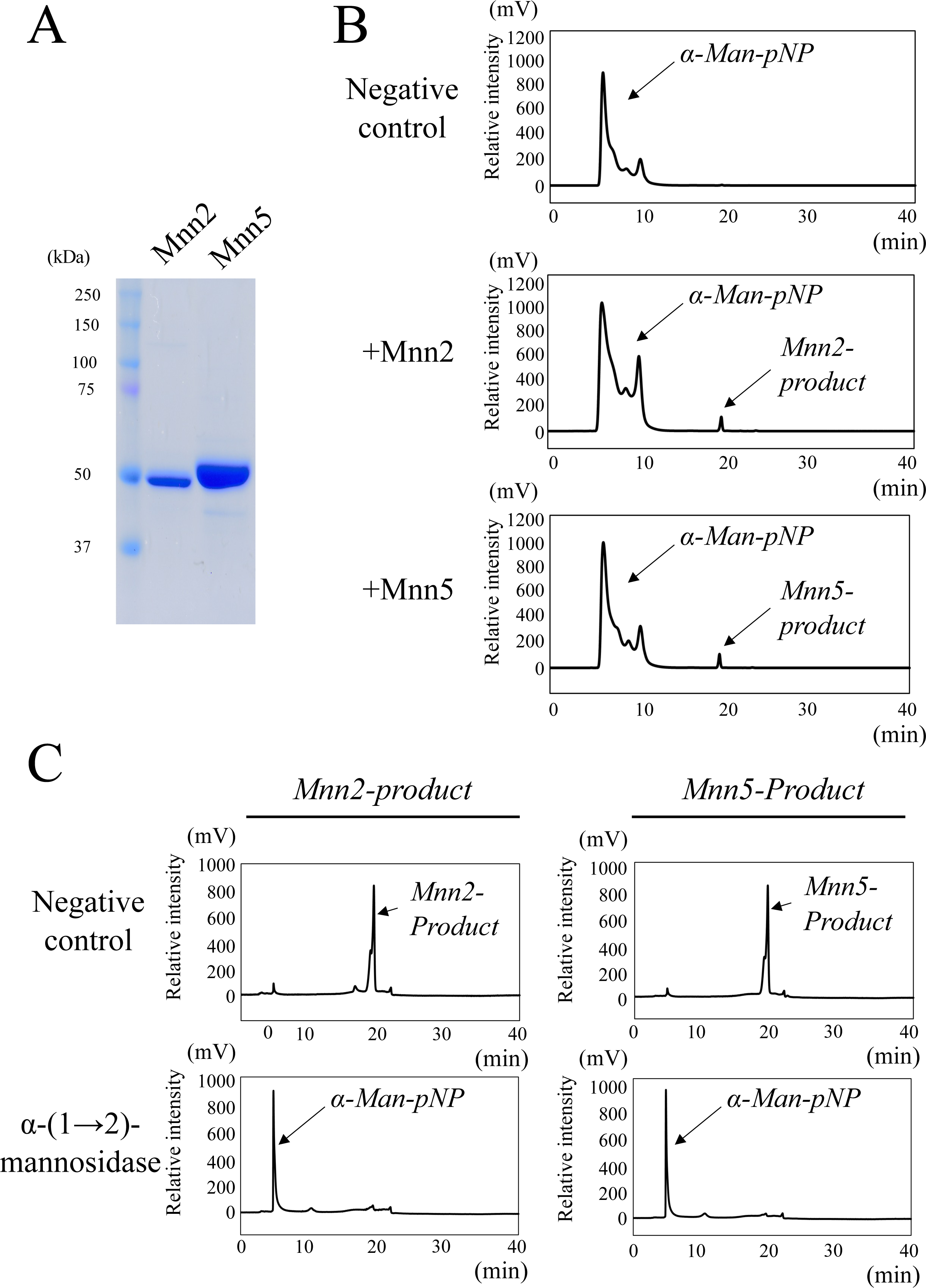
Mannosyltransferase activities of Mnn2 and Mnn5 *in vitro*. (A) SDS-PAGE analysis of purified recombinant Mnn2 and Mnn5. The purified recombinant proteins were separated by SDS-PAGE using a 5–20% gradient polyacrylamide gel. (B) Chromatograms of Mnn2 and Mnn5 mannosyltransferase activity assays using p-nitrophenyl α-D-mannopyranoside as the acceptor substrate. The reaction mixture containing 50 mM HEPES-NaOH (pH 6.8), 100 mM NaCl, 30 mM KCl, 5% glycerol, 1 mM MnCl_2_, 1.5 mM α-Man-pNP (acceptor substrate), 5 mM GDP-Man (donor substrate), and 6.0 µg of purified Mnn2 and Mnn5 was incubated at 30°C for 16 h, respectively. Chromatograms show typical results of the assay without enzyme (negative control, *upper panel*), with Mnn2 (*middle panel*), and with Mnn5 (*lower panel*). The assay without enzyme yielded only peaks derived from the α-Man-pNP at 5-10 min, whereas fractions with Mnn2 and Mnn5 displayed reaction products (termed Mnn2-product and Mnn5-product) at 19.0 min, respectively. (C) Structural analysis of Mnn2-product (at left) and Mnn5-product (at right) using α-1,2-mannosidase. Upper panels show chromatographs of the purified Mnn2-product or Mnn5-product. Purified Mnn2-product and Mnn5-product were reacted with α-1,2-mannosidase (lower panels). Both Mnn2-product and Mnn5-product could be reacted with α-1,2-mannosidase and digested to α-Man-pNP.

### Colony morphology of A1151, Δ*mnn2*, Δ*mnn5*, and Δ*mnn2*Δ*mnn5*

To uncover the physiological roles of Mnn2 and Mnn5, we generated single gene disruptants for *mnn2* and *mnn5*, as well as a double disruptant for *mnn2*/*mnn5* in *A. fumigatus* (Fig. S1). Subsequently, we compared colony morphology at 37°C. The Δ*mnn2* and Δ*mnn5* strains exhibited growth patterns similar to the A1151 strain. However, intriguingly, the Δ*mnn2*Δ*mnn5* strain displayed aberrant colony morphology and showed a reduction in conidiophore formation (Fig. 2A). Consequently, our investigation extended to the evaluation of mycelial elongation rate and the number of mycelia per colony area for each strain (Fig. 2B and C).

**Fig. 2.**
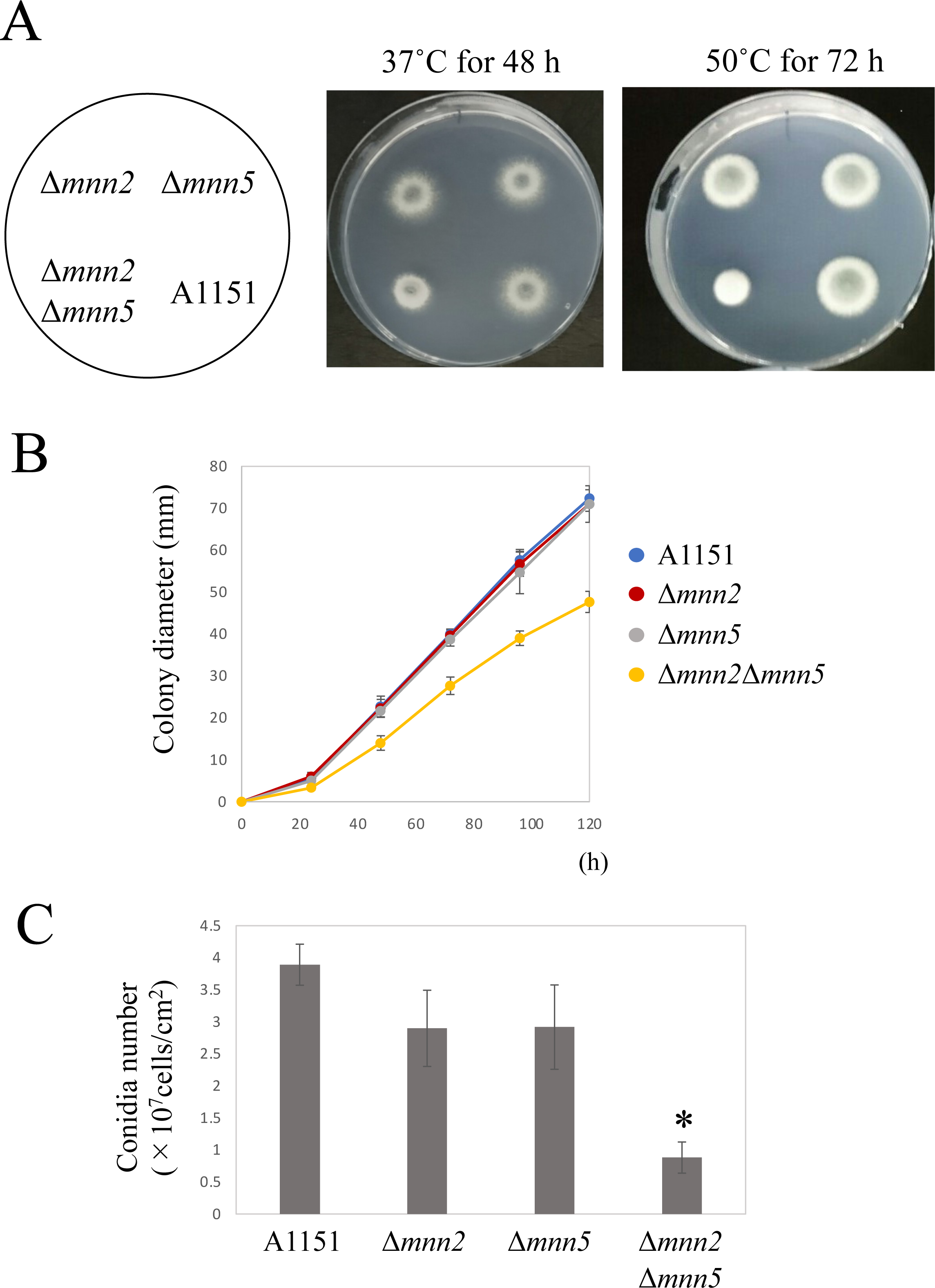
Phenotype of *mnn2* and *mnn5* single or double disruptants. (A) Colony morphology of A1151, Δ*mnn2*, Δ*mnn5*, and Δ*mnn2* Δ*mnn5* on MM agar at 37 for 2 days and 50°C for 3 days, respectively. The agar medium was inoculated with 1.0 × 10^4^ conidiospores. (B) Colony diameter of A1151, Δ*mnn2*, Δ*mnn5*, and Δ*mnn2* Δ*mnn5* on MM agar at 37°C for 0, 24, 48, 72, 96, 120 hours, respectively. (C) Conidia number per colony area of A1151, Δ*mnn2*, Δ*mnn5*, and Δ*mnn2* Δ*mnn5* on MM agar at 37°C for 5 days. Asterisks indicate significant differences (*, P < 0.05; Welch’s t test) from the results for the A1151 strain.

Significantly, the diameters of the Δ*mnn2*Δ*mnn5* colonies were 0.59-, 0.62-, 0.69-, 0.68-, and 0.66-fold smaller than those of the A1151 colonies when cultured on MM at 37°C for 24, 48, 72, 96, and 122 h, respectively (Fig. 2B). Furthermore, the number of conidia per colony area in Δ*mnn2*Δ*mnn5* was 0.23-fold lower than that in the A1151 strain. These compelling findings strongly indicate that Mnn2 and Mnn5 possess overlapping functions and play crucial roles in the processes of hyphal elongation and conidia formation in *A. fumigatus*.

### Structural analysis of mannoprotein and FTGM from A1151, Δ*mnn2*, Δ*mnn5*, and Δ*mnn2*Δ*mnn5*

The study investigated the functions of Mnn2 and Mnn5 *in vivo* by analyzing total mannoproteins and GMs from A1151, Δ*mnn2*, Δ*mnn5*, and Δ*mnn2*Δ*mnn5* strains. The mannoproteins and GMs were extracted from mycelia using autoclaving at 121°C for 2 h with 100 mM citrate buffer (pH 7.0). They were then treated with 0.1 M HCl to remove the β-(1→5)-/β-(1→6)-galactofuran side chain, and O-linked glycans such as OMGM were removed using the β-elimination method. The N-glycans and FTGM α-core-mannan structure were analyzed using ^1^H-NMR spectroscopy. The common four signals (A-D) of A1151, Δ*mnn2*, Δ*mnn5*, and Δ*mnn2*Δ*mnn5* strains emerged, indicating the H-1 signal of the chemical shift of the α-(1→2)-/α-(1→6)-linked mannan of FTGM (Fig. 3A) (Kudoh et al. 2015). However, the 4.9 ppm signal, which indicates the H-1 signal of the chemical shift of the α-(1→6)-linked mannan, was specifically detected in the Δ*mnn2*Δ*mnn5* strain (Fig. 3A). The one-dimensional total correlation spectroscopys (TOCSY) spectrum of Δ*mnn2*Δ*mnn5,* recorded by the irradiation of the signal at 4.9 ppm corresponding to the branching α-(1→6)-mannosyl residue, was very similar to the previously reported spectrum of long-length α-(1→6)-linked mannan from *S. cerevisiae* Δ*mnn2* (Shibata et al. 1996). These results suggest that the double disruption of *mnn2* and *mnn5* resulted in the α-(1→6)-linked mannan backbone becoming exposed and detectable in *A. fumigatus*.

**Fig. 3.**
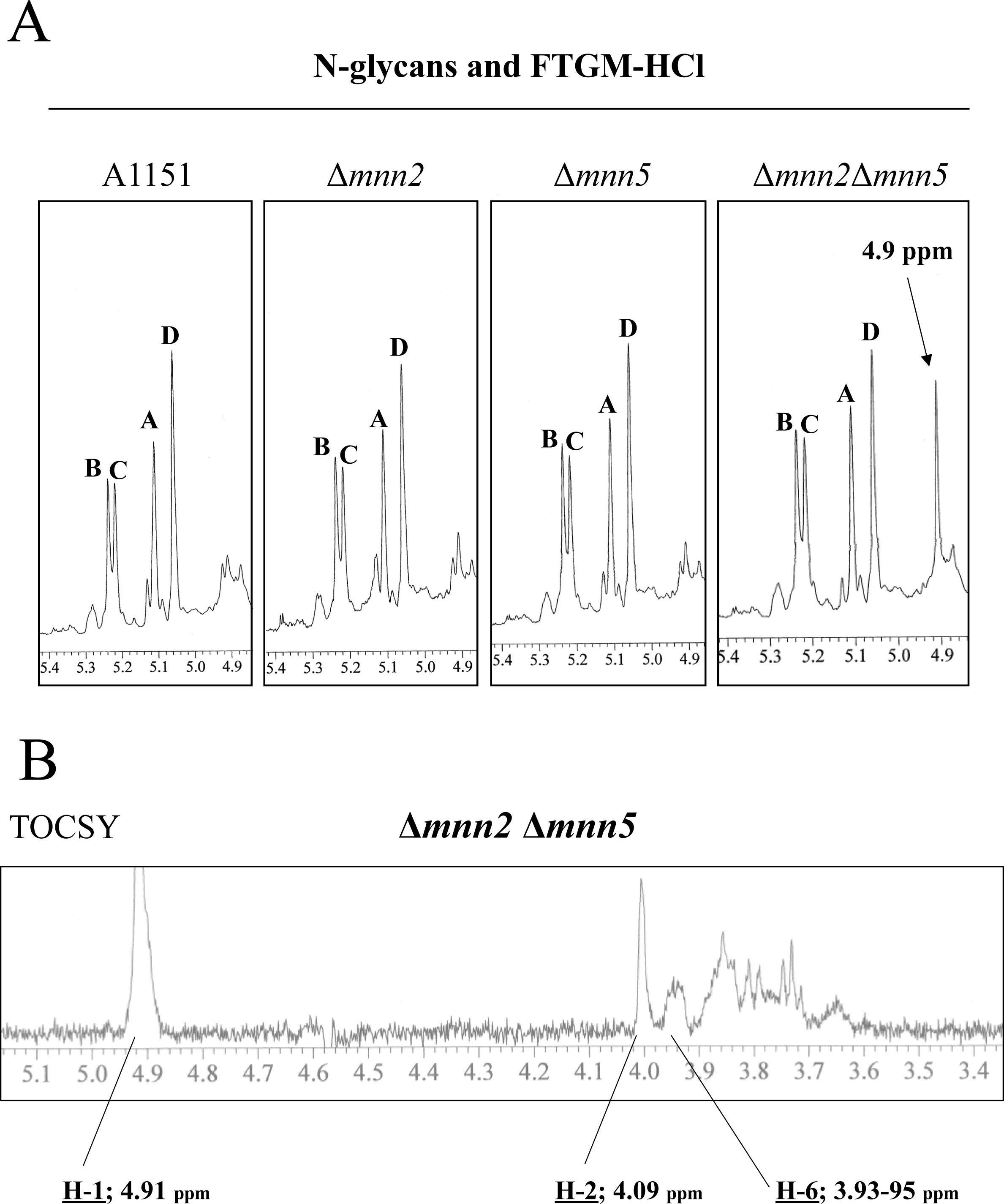
^1^H-NMR analysis of N-glycan and core-mannan of FTGM with removedgalactofuran side chains (FTGM-HCl) from WT (A1151), Δ*mnn2,* Δ*mnn5,* and Δ*mnn2*Δ*mnn5* strains. The signals A (at 5.104 ppm), B (5.233), C (5.216), and D (5.054) of the ^1^ H-NMR spectra are derived from H-1 at the C-1 position of the underlined Man residues in the structures –(1→6)-α-<u>Man</u>-(1→2)-α-Man-(1→2)-α-Man-(1→2)-α-Man-(1→6)-(A), –(1→6)-α-Man-(1→2)-α-<u>Man</u>-(1→2)-α-Man-(1→2)-α-Man-(1→6)-(B), – (1→6)-α-Man-(1→2)-α-Man-(1→2)-α-<u>Man</u>-(1→2)-α-Man-(1→6)-(C) and –(1→6)-αMan-(1→2)-α-Man-(1→2)-α-Man-(1→2)-α-<u>Man</u>-(1→6)-(D). The signals at 4.9 ppm in the Δ*mnn2*Δ*mnn5* of the ^1^ H-NMR spectra are from H-1 at the C-1 position of the underlined Man residue in the structures t-α-<u>Man</u>-(1→6)-α-Man and –(1→6)-α-<u>Man</u>-(1→6)-. The proton chemical shifs were referenced relative to internal acetone at δ 2.225 ppm.

### Expression of Mnn2 and Mnn5 in *S. cerevisiae* Δ*Scmnn2*Δ*Scmnn5*

To assess whether Mnn2 and Mnn5 can complement the function of *Sc*Mnn2p and *Sc*Mnn5p, we developed a heterologous expression strain in *S. cerevisiae* with a Δ*Scmnn2*Δ*Scmnn5* genetic background. The outer chain structure of BY4741, Δ*Scmnn2*, Δ*Scmnn5,* Δ*Scmnn2,* Δ*Scmnn5*, Δ*Scmnn2,* Δ*Scmnn5* expressing *mnn2*, and Δ*Scmnn2,* Δ*Scmnn5* expressing *mnn5* strains was analyzed using ^1^H-NMR (Fig. 4). The chemical shift around 5.0–5.3 ppm observed in the wild-type BY4741 budding yeast indicates the presence of the α-(1→2)-and α-(1→3)-Man side chain of the N-glycan outer chain (Tsai et al. 1984). In the Δ*Scmnn5* strain, chemical shifts appeared at 5.04 ppm and 5.14 ppm, indicating the presence of Man-α-(1→2)-Man and Man-α-(1→2)-Man-α-(1→6)-Man, respectively. Additionally, a chemical shift of 4.9 ppm was observed in the Δ*Scmnn2* strain, indicating the presence of α-(1→6)-linked mannan (Fig. 4). Heterologous expression of Mnn2 and Mnn5 in Δ*Scmnn2*Δ*Scmnn5*, respectively, resulted in the appearance of chemical shifts at 5.04 and 5.14 ppm, although the chemical shift at 4.9 ppm was not completely eliminated (Fig. 4). These results indicate that Mnn2 and Mnn5 partially complement the function of budding yeast *Sc*Mnn2p. This is consistent with the appearance of a chemical shift of 4.9 ppm for the first time in *A. fumigatus* by the double disruption of *mnn2* and *mnn5*.

**Fig. 4.**
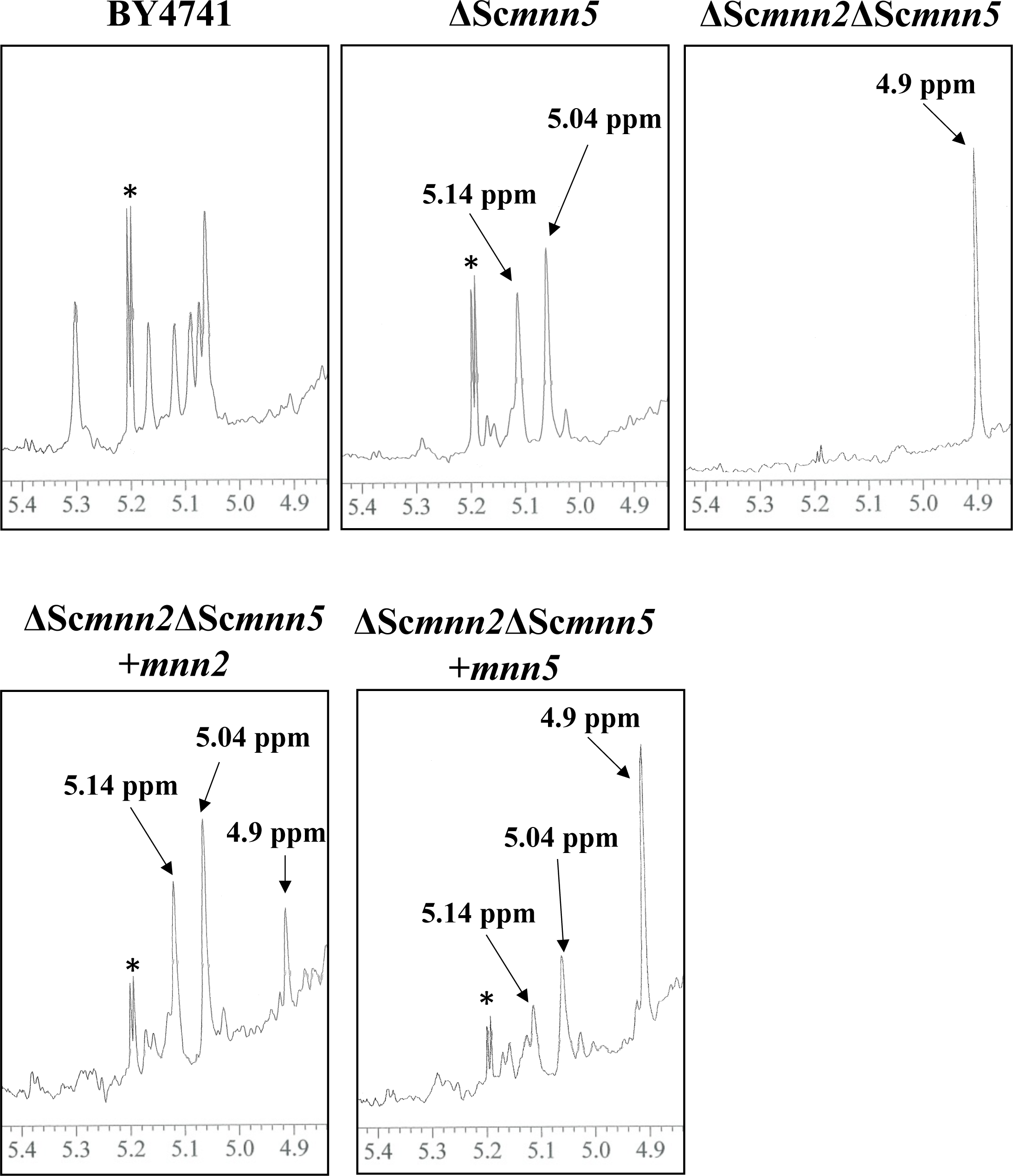
^1^H-NMR analysis of N-glycan outer chains from yeast WT (BY4741), ΔSc*mnn5*, ΔSc*mnn2*ΔSc*mnn5*, ΔSc*mnn2*ΔSc*mnn5* + *mnn2*, and ΔSc*mnn2*ΔSc*mnn5* + *mnn5* Strains. The 1H-NMR spectra depict the N-glycan outer chains of various yeast strains. In the WT strain (BY4741), signals at 5.0∼5.3 ppm correspond to H-1 at the C-1 position of Man residues in the α-(1→2)-Man and α-(1→3)-Man side chain of the outer chain. In *ΔScmnn5*, signals at 5.04 and 5.14 ppm indicate H-1 at the C-1 position of Man in the structures t-α-<u>Man</u>-(1→2)-α-Man and Man-(1→2)-α-<u>Man</u>-(1→6)-α-Man. In *ΔScmnn2ΔScmnn5*, signals at 4.9 ppm are from H-1 at the C-1 position of the underlined Man residue in the structures t-α-<u>Man</u>-(1→6)-α-Man and –(1→6)-α-<u>Man</u>-(1→6)-. Asterisks denote unidentified NMR signal. Proton chemical shifs are referenced relative to internal acetone at δ 2.225 ppm.

### Structural analysis of mannoprotein and FTGM from A1151, Δ*mnn2*Δ*mnn5*, Δ*mnn2*Δ*mnn5*Δ*mnn9*, Δ*mnn2*Δ*mnn5*Δ*van1*, and Δ*mnn2*Δ*mnn5*Δ*anpA*

In order to identify the glycosyltransferases involved in the biosynthesis of α-(1→6)-linked mannan in *A. fumigatus*, mannoproteins from A1151, Δ*mnn2*Δ*mnn5*, Δ*mnn2*Δ*mnn5*Δ*mnn9*, Δ*mnn2*Δ*mnn5*Δ*van1,* and Δ*mnn2*Δ*mnn5*Δ*anpA* were extracted and their structures were analyzed (Fig. 5). The disruption of *mnn9* or *van1* in the Δ*mnn2*Δ*mnn5* strain resulted in the disappearance of the 4.9 ppm chemical shift observed in Δ*mnn2*Δ*mnn5* (Fig. 5). This suggests that Mnn9 and Van1 are essential for the biosynthesis of α-(1→6)-linked mannan in *A. fumigatus*. In the Δ*mnn2*Δ*mnn5*Δ*anpA* strain, the signal of FTGM was absent, but the signal of α-(1→6)-linked mannan was clearly detected. This indicates that AnpA is involved only in the biosynthesis of FTGM, but not α-(1→6)-linked mannan (Fig. 5).

**Fig. 5.**
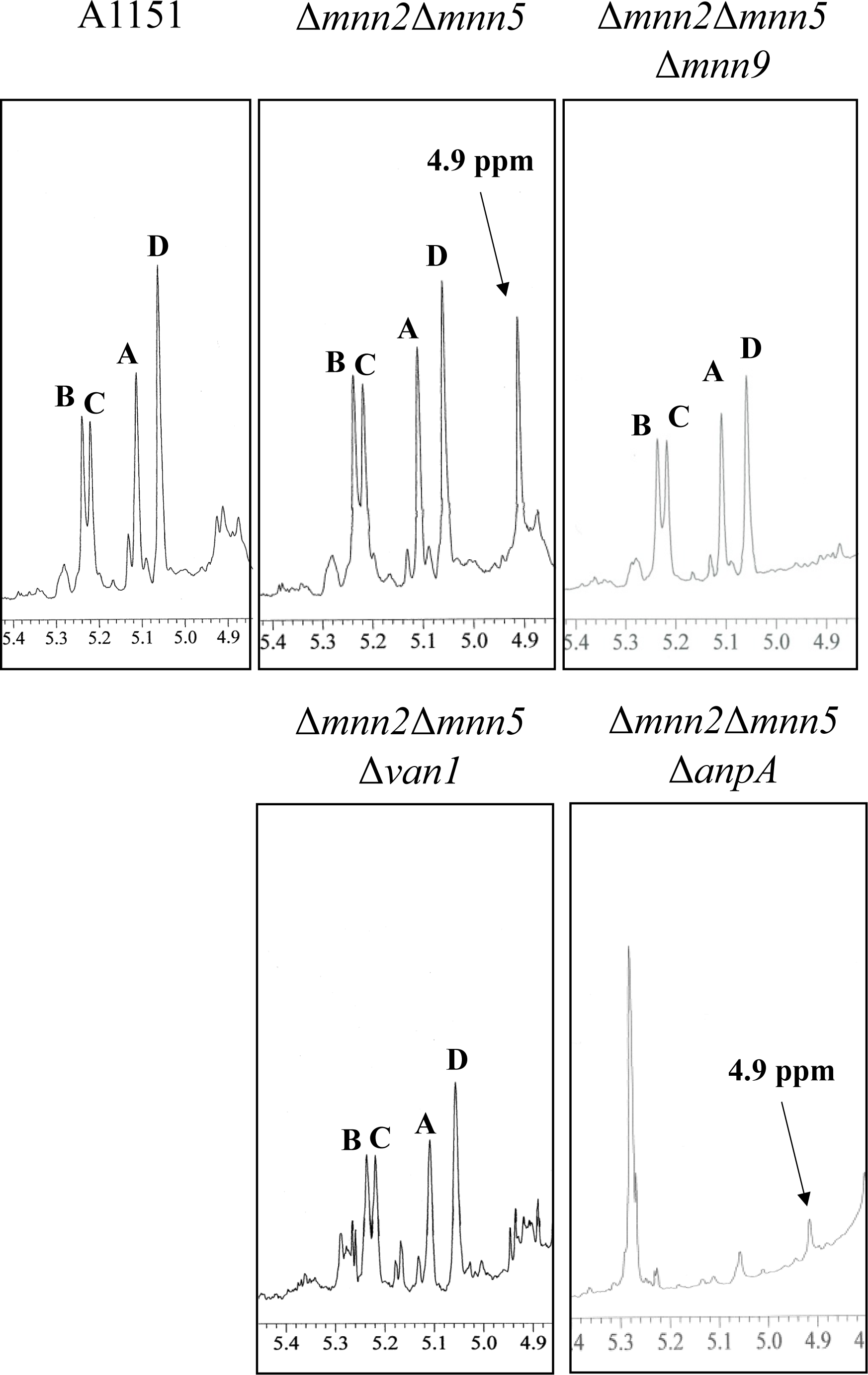
^1^H-NMR analysis of N-glycan and core-mannan of FTGM with removed galactofuranside chains (FTGM-HCl) from WT (A1151), Δ*mnn2*Δ*mnn5*, Δ*mnn2*Δ*mnn5*Δ*mnn9*, Δmnn2Δmnn5Δ*van1*, and Δ*mnn2*Δ*mnn5*Δ*anpA* strains strains. The Signals A (at 5.104 ppm), B (5.233), C (5.216), and D (5.054) of the ^1^ H-NMR spectra are derived from H-1 at the C-1 position of the underlined Man residues in the structures-(1→6)-α-<u>Man</u>-(1→2)-α-Man-(1→2)-α-Man-(1→2)-α-Man-(1→6)-(A), –(1→6)-α-Man-(1→2)-α-<u>Man</u>-(1→2)-α-Man-(1→2)-α-Man-(1→6)-(B), –(1→6)-α-Man-(1→2)-α-Man-(1→2)-α-<u>Man</u>-(1→2)-α-Man-(1→6)-(C) and –(1→6)-αMan-(1→2)-α-Man-(1→2)-α-Man-(1→2)-α-<u>Man</u>-(1→6)-(D). The signals at 4.9 ppm in the Δ*mnn2*Δ*mnn5* of the ^1^ H-NMR spectra are from H-1 at the C-1 position of the underlined Man residue in the structures t-α-<u>Man</u>-(1→6)-α-Man and –(1→6)-α-<u>Man</u>-(1→6)-. Asterisks indicate unidentified NMR signals. Proton chemical shifs are referenced relative to internal acetone at δ 2.225 ppm.

### Mannosyltransferase activities of Mnn9 and Van1 *in vitro*

To explore the mannosyltransferase activities of Mnn9 (AFUB_018530) and Van1 (AFUB_031580) from *A. fumigatus*, we generated individual N-terminal 6×Histagged recombinant proteins using a bacterial expression system. Mnn9 and Van1 were expressed without the presumed transmembrane domains, covering amino acid residues 1-35 and 1-69, respectively. While Mnn9 yielded solubilized protein successfully, unfortunately, Van1 proved to be insoluble (data not shown). In an attempt to address this, we removed the region from amino acid numbers 387 to 499, anticipating no impact on the function of the Van1 C-terminus. This modification led to the successful procurement of solubilized Van1 protein. The obtained recombinant proteins underwent sodium dodecyl sulfate-polyacrylamide gel electrophoresis (SDS-PAGE) (Fig. 6). Despite detecting mannosyltransferase activities, Mnn9 and Van1 exhibited no activity when tested individually. This suggests that neither of them functions as a glycosyltransferase in homodimers (Fig. 7). Subsequently, we co-expressed 6×Histagged Mnn9 and untagged Van1. Purification with Ni-agarose yielded Van1 along with Mnn9, even without the 6xHIS tag on Van1 (Fig. 6). This indicated the formation of a protein complex between Mnn9 and Van1. Interestingly, multiple peaks, potentially representing mannosyl polymers, were detected under the Mnn9 and Van1 co-expression conditions (Fig. 7). This peak was susceptible to digestion by jack bean mannosidase (α-(1→2,3,6)-mannosidase) but not by α-(1→2,3)-mannosidase, conclusively confirming it as an α-(1→6)-mannose polymer (Fig. 8). These results elucidate that Mnn9 and Van1 exclusively function as mannan polymerases when they form heterodimers.

**Fig. 6.**
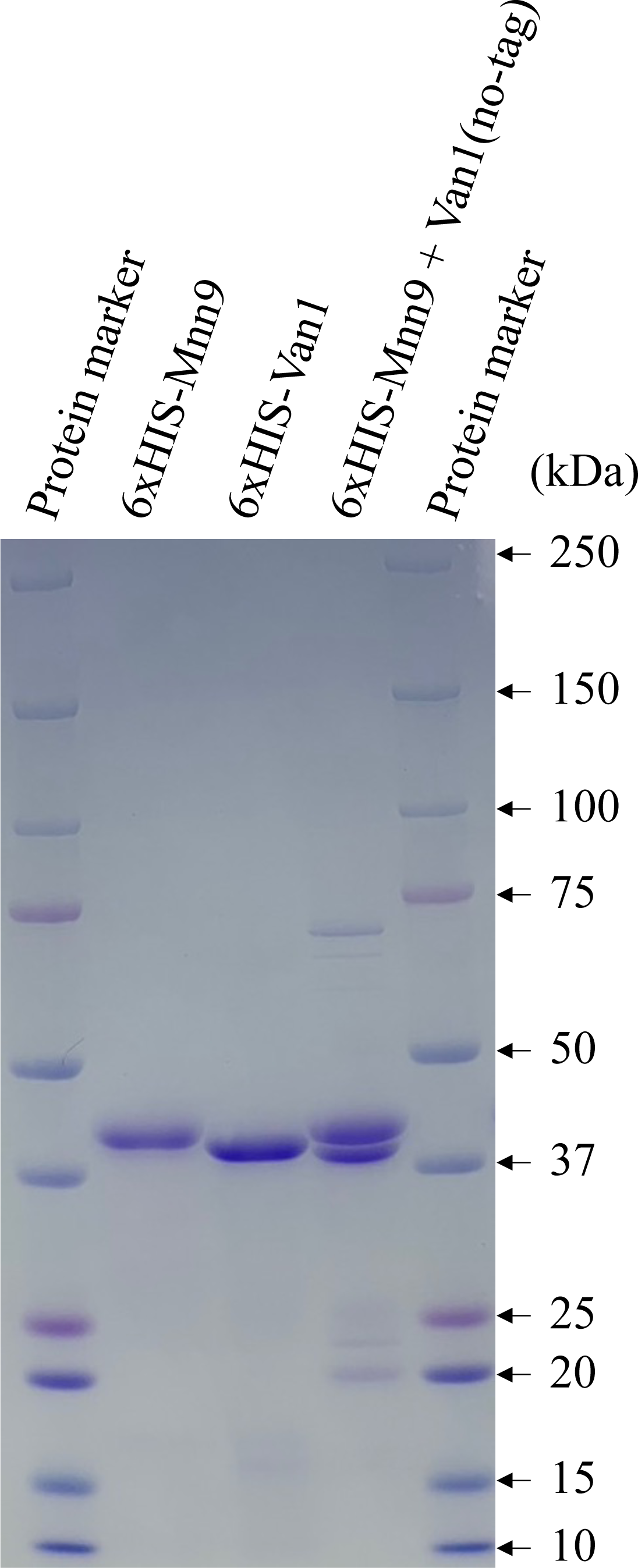
Mnn9 and Van1 of *Aspergillus fumigatus* form heterodimers *in vitro*. SDS-PAGE analysis of purified recombinant 6×His-tagged Mnn9, 6×His-tagged Van1, and 6×His-tagged Mnn9 + non-tagged Van1. Lanes 2-4 show 3 μg of purified proteins (Lane 1: 6×His-tagged Mnn9, Lane 2: 6×His-tagged Van1, Lane 3: 6×His-tagged Mnn9 + non-tagged Van1 while Lane 1 and 5 display 5 μL of Precision Plus Protein™ Dual Color Standard (Bio-Rad).

**Fig. 7.**
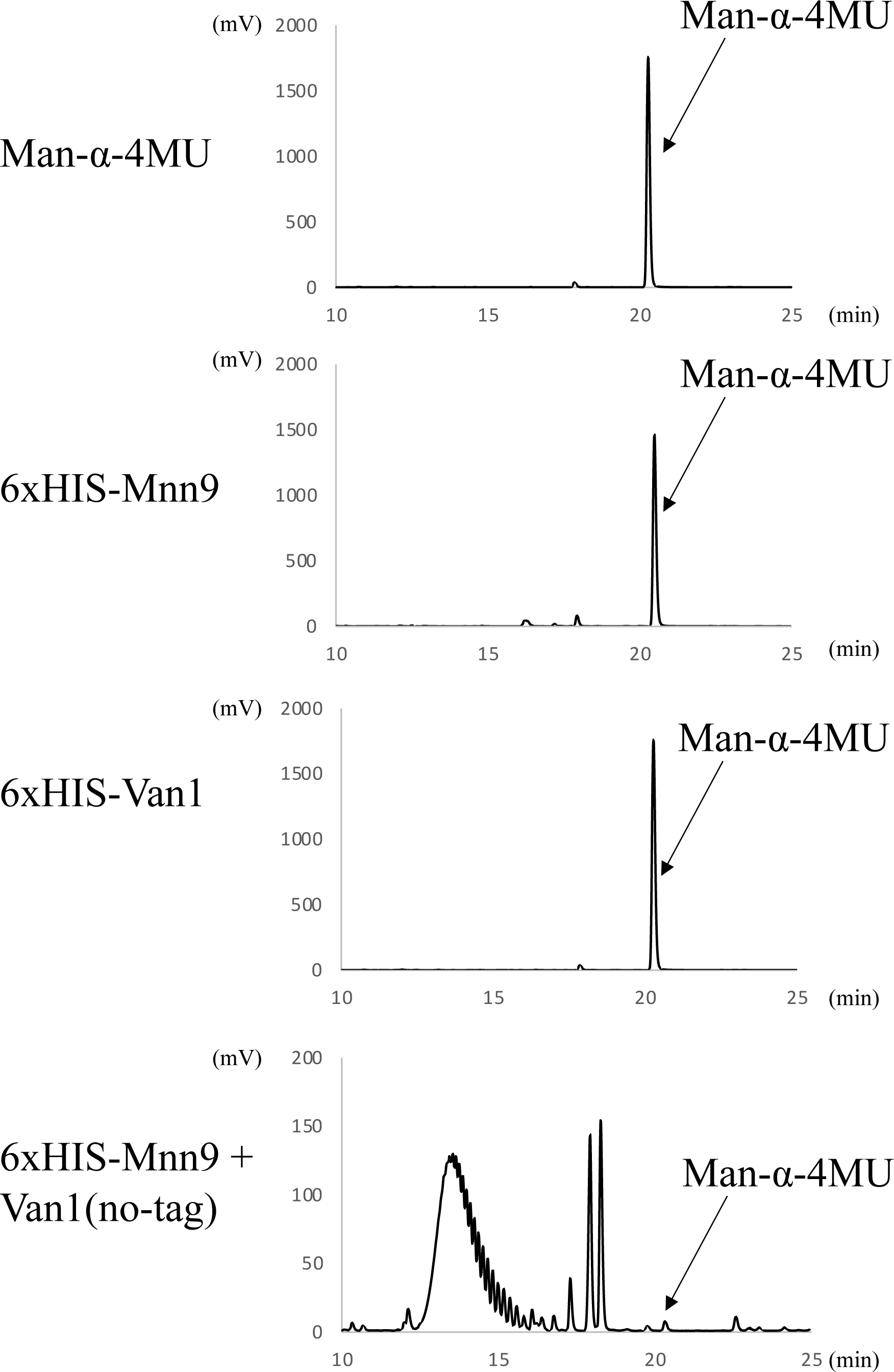
Mannan polymerase activity of heterodimeric Mnn9 and Van1 *in vitro*. Chromatograms of Mnn9 and Van1 mannosyltransferase activity assays using 4-methylumbelliferyl α-D-mannopyranoside as the artificial acceptor substrate. The assays were conducted in a reaction mixture (40 µl) containing 50 mM HEPES-NaOH (pH 6.8), 100 mM NaCl, 30 mM KCl, 5% glycerol, 1 mM MnCl_2_, 1.5mM α-Man-4MU (acceptor substrate), 5 mM GDP-Man (donor substrate), and 10 µg of purified enzymes at 30°C for 16 h. Chromatograms depict results of the assay without enzyme (negative control) and with Mnn9, Van1, and Mnn9 + Van1, respectively. Without enzyme and Mnn9 alone and Van1 alone assays show peaks derived from Man-α-4MU at 22.0 min, whereas fractions with Mnn9 + Van1 display reaction products.

**Fig. 8.**
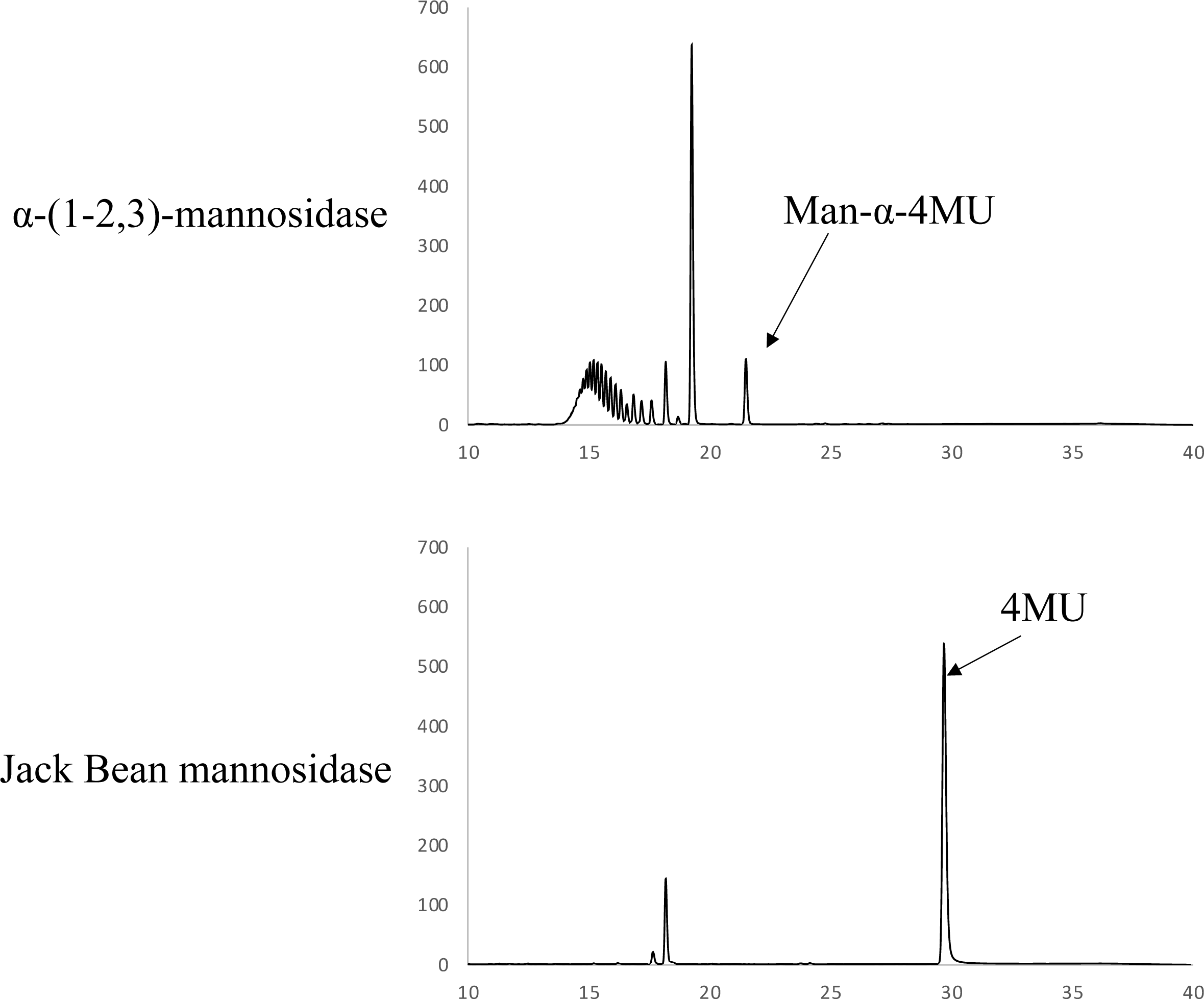
Heterodimer of Mnn9 and Van1 synthesized α-(1→6)-mannan *in vitro*. Structural analysis of Mnn9+Van1 products using α-(1 →2,3)-mannosidase and Jack Bean mannosidase. Purified Mnn9 + Van1 product was reacted with α-(1 →2,3) –mannosidase (upper panel) and Jack Bean mannosidase (lower panel). Both Mnn9 + Van1 product could be reacted with Jack Bean mannosidase and digested to 4MU.

## Discussion

In this investigation, we delve into the functional roles of Mnn2 and Mnn5, critical contributors to hyphal growth and conidial formation through the biosynthesis of mannan structures in *A. fumigatus*. Unlike β-glucan and chitin, Mannan structures in yeast and filamentous fungi are not indispensable for growth but play a crucial role in maintaining overall cell wall integrity.

Our ^1^H-NMR analysis of mannoproteins unveiled the presence of outer chain like mannan structures in *A. fumigatus* mycelium. The distinctive chemical shift originating from the α-(1→6)-linked mannan core chain becomes discernible when the side chain is absent (Raschke et al. 1973). Both Mnn2 and Mnn5 emerge as pivotal catalysts in side-chain biosynthesis, indicating their collaborative role in transferring the initial α-(1→2)-mannosyl residues to the α-(1→6)-linked mannan backbone (Fig. 9). In *S. cerevisiae*, Mnn5p is recognized for transferring the second α-(1→2)-mannosyl residue of the side chain (Rayner and Munro. 1998). However, upon heterologous expression of Mnn2 and Mnn5, the mannan structure of Δ*Scmnn2*Δ*Scmnn5* was restored to that of Δ*Scmnn5* (Fig. 4), implying the functional homology of Mnn2 and Mnn5 with Mnn2p. This observation suggests that the chain length of the α-(1→2)-mannosyl side chain of α-(1→6)-linked mannan in *A. fumigatus* is only one. In other words, both Mnn2 and Mnn5 are responsible for the biosynthesis of the first mannosyl residues of α-(1→6)-linked mannan. Moving forward, we uncovered the involvement of Mnn9 and Van1, members of the GT62 family, in mycelial α-(1→6)-linked mannan biosynthesis (Fig. 5). The heterodimer of Mnn9 and Van1 was found to function as a mannan polymerase (Fig. 6, 7, 8). Contrary to prior investigations documenting Mnn9’s manifestation of mannosyltransferase activity in isolation (Henry, 2016), our findings reveal that neither Mnn9 nor Van1 operates independently (Fig. 7). This outcome contrasts with the singular activity of the recombinant AnpA involved in the biosynthesis of fungal-type galactomannan (Kadooka et al. 2022b). Furthermore, our results align with the obliteration of the α-(1→6)-linked mannan-derived chemical shift following the singular disruption of either *mnn9* or *van1* (Fig. 5). These findings underscore the essential collaborative role of both *mnn9* and *van1* in the biosynthesis of the outer chain like mannan in *A. fumigatus*. In the context of *A. fumigatus*, the *mnn2* and *mnn5* double-disruption strain displayed a reduced mycelial elongation rate and diminished ability to form conidiophores (Fig. 2). Notably, reports indicate that Δ*mnn9* and Δ*van1* do not exhibit the Δ*mnn2*Δ*mnn5*-like phenotype (Henrry et al. 2016, Kadooka et al. 2022b). This discrepancy suggests that Mnn2 and Mnn5 may also be involved in the biosynthesis of mannans beyond the outer chain. Additionally, a strong correlation between outer chain structure and MIPC has been noted in *S. cerevisiae* (Morimoto and Tani. 2015). The lack of growth retardation in *A. fumigatus* upon the loss of ceramides such as GIPCs implies a compensatory relationship between these structures in their function (Kotz et al. 2010, Engel et al. 2015, Seegar et al. 2022). The intricate landscape of mannan biosynthesis in *A. fumigatus* remains complex and warrants further elucidation in future studies.

**Fig. 9.**
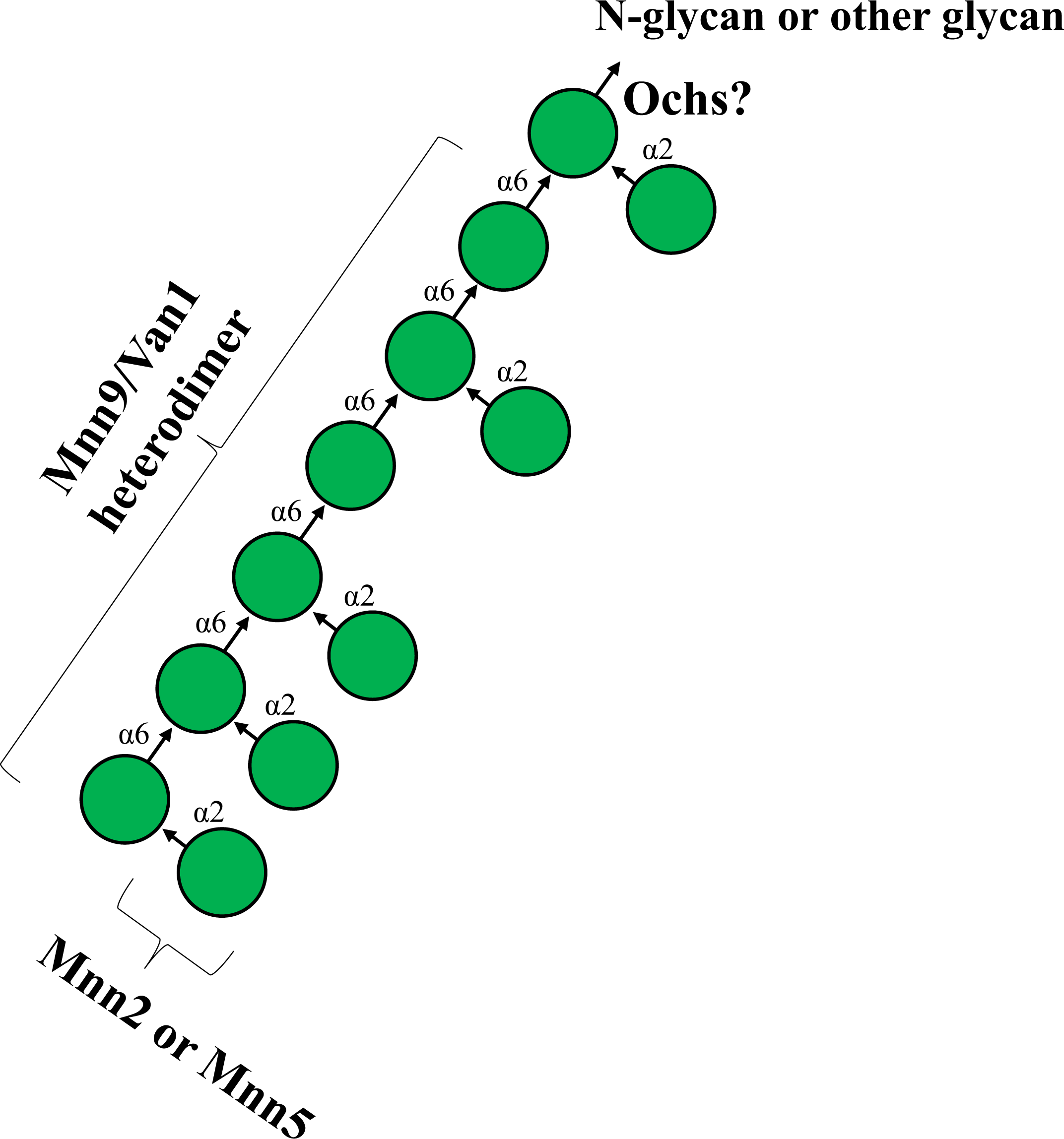
Summary Model of N-Glycan Outer Chain Mannan Biosynthesis in *Aspergillus fumigatus*. Ovals represent the initial specific α-(1→6)-mannosyltransferase. Mnn9/Van1 heterodimer represents α-(1→6)-mannan polymerase. Mnn2 and Mnn5 represent α-(1→2)-mannosyltransferase.

Mnn9 and Van1 in *A. fumigatus* have undergone extensive scrutiny, with recent findings linking them to the biosynthesis of the conidia-specific G3Man structure (Liu et al 2023). Our study contributes a novel revelation by showcasing, for the first time, the presence of α-(1→6)-linked mannan in both mycelia and conidia. Intriguingly, Mnn9 and Van1 play a role in the biosynthesis of α-(1→6)-linked mannan in mycelial and conidial forms (Liu et al. 2023). However, the structural disparities in the side chains between conidial and mycelial outer chain like mannans suggest that, in comparison to the complex G3Man, the mycelial outer chain like mannan exhibits a simpler structure (Fig. 9). This structural distinction is likely attributed to the necessity for a complex structure, such as G3Man, in conidia due to the high percentage of β-glucan and the absence of galactosaminogalactan (GAG) (Liu et al. 2023). Conversely, the presence of α-glucan and GAG in the mycelium may render a simpler mannan structure sufficient. The presence of mannans with different side-chain structures in the cell walls of both conidia and mycelia remains unparalleled among known polysaccharides. Investigating the specific functional roles of these mannans in conidia and mycelia presents an intellectually compelling avenue for research. The physiological significance of these mannan structures warrants further elucidation.

Notably, while the loss of outer chains in budding and fission yeasts significantly decreases the molecular weight of invertase and acid phosphatase (Nakayama et al 1992, Ohashi et al 2020), a similar phenomenon was not observed in *A. fumigatus* (Du et al 2020). This discrepancy may be attributed to Gel1 not having an outer chain N-glycan or having shorter chains compared to yeast. The observation aligns with the fact that the loss of outer chain structure causes substantial growth defects in yeast but does not result in a remarkable phenotype in *A. fumigatus* (Nakayama et al 1992, Ohashi et al 2020, Henrry et al. 2016, Kadooka et al. 2022a). Du et al. proposed that α-mannan elongation occurs not via N-glycan but through O-linked mannose in cell wall mannoproteins (Du et al 2020). Unlike yeasts, which heavily rely on outer chain structures for cell wall integrity, the presence of other mannan structures, such as FTGM, in *A. fumigatus* may diminish the importance of outer chain structures. Conversely, certain species, such as *N. crassa,* emphasize the significance of outer chain structure even in filamentous fungi, showcasing the intriguing diversity of mannan structures, including the outer chain, within the phylum Ascomycota (Maddi et al 2010).

In summary, our investigations into Mnn2 and Mnn5 have unveiled the presence of an outer chain like mannan structure in the mycelia of *A. fumigatus*. These findings mark a substantial contribution to our comprehension of the structure and biosynthesis of filamentous fungal cell wall, with potential implications for the development of antifungal and agrochemical agents.

## Materials and Methods

### Strains and medium

The *A. fumigatus* strains utilized in this investigation are detailed in Supplemental Table 1 (Table S1), with *A. fumigatus* A1160 and A1151 strains serving as the parental and control strains, respectively.

Cultures were developed on *Aspergillus* minimal medium (1% [wt/vol] glucose, 0.6% [wt/vol] NaNO_3_, 0.052% [wt/vol] KCl, 0.052% [wt/vol] MgSO_4_·7H_2_O, and 0.152% [wt/vol] KH_2_PO_4_, supplemented with Hutner’s trace elements [pH 6.5]). For the A1160 strain cultivation, 1.22 g/l and 1.21 g/l uracil and uridine were incorporated into the MM. The *S. cerevisiae* strains employed in this study are enumerated in Table S1. BY4741 and Δ*mnn5* strains were procured from the Yeast Knockout (YKO) Collection (Horizon Discovery Ltd., UK). Growth for these strains occurred on YPD medium or synthetic complete (SC) medium.

### Construction of *mnn2* and *mnn5* gene disruption strains

The disruption of the *mnn2* and *mnn5* genes in *A. fumigatus* A1160 was achieved through *AnpyrG* insertion. A gene replacement cassette, consisting of the homology arm at the 5′ end of the *mnn2* and *mnn5* genes and the homology arm at the 3′ end of the *mnn2* and *mnn5* genes, was generated by recombinant PCR. The *A. fumigatus* A1151 genomic DNA served as the template, and the primer pairs xxxx-1/xxxx-2, and xxxx-3/xxxx-4 were used for amplification (where “xxxx” indicates *mnn2* or *mnn5*) (Table S2). Simultaneously, the *AnpyrG* marker was amplified by recombinant PCR using pHSG396-AnpyrG (Kadooka et al. 2022b) as the template and the primer pair pHSG396-F/pHSG396-R. The resulting DNA fragment, amplified with primers xxxx-1 and xxxx-4, was employed for the transformation of *A. fumigatus* A1160, yielding Δ*mnn2* and Δ*mnn5* strains. MM agar plates lacking uracil and uridine were utilized for the selection of transformants. The successful introduction of *AnpyrG* into each gene locus was confirmed through PCR, using the primer pairs mnn2-F/pyrG-R and pyrG-F/mnn2-R, as well as mnn5-F/pyrG-R and pyrG-F/mnn5-R, respectively (Fig. S1).

### Construction of pHSG396-ptrA plasmid

The pyrithiamine resistance gene (*ptrA*) was amplified by PCR, utilizing the plasmid pPTR I (TAKARA, Japan) as a template and the primer pairs pHSG396-ptrA-IF-F and pHSG396-ptrA-IF-R. Subsequently, the amplified fragment was incorporated into the *Bam* HI site of pHSG396 using the In-Fusion HD cloning kit (TAKARA, Japan), resulting in the generation of pHSG396-ptrA.

### Construction of *mnn2* and *mnn5* double-disruption strains

The *mnn5* gene was disrupted in *A. fumigatus* Δ*mnn2* by *ptrA* or *hph* insertion. A gene replacement cassette, including the homology arm at the 5′ end of the *mnn5* gene and the homology arm at the 3′ end of the *mnn5* gene,was generated by recombinant PCR using *A. fumigatus* A1151 genomic DNA as the template and the primer pairs mnn5-1/mnn5-2 and mnn5-3/mnn5-4, respectively (Table S2). The *ptrA* and *hph* markers were amplified by recombinant PCR using pHSG396-ptrA and pHSG396-hph (Kadooka et al. 2022a) as templates and the primer pair pHSG396-F/pHSG396-R. The resulting DNA fragments, amplified with primers mnn5-1 and mnn5-4, were used to transform *A. fumigatus* Δ*mnn2*, generating Δ*mnn2*Δ*mnn5* (*ptrA*) and Δ*mnn2*Δ*mnn5* (*hph*), respectively. MM agar plates supplemented with 0.1 µl/ml pyrithiamine or 200 µl/ml hygromycin B were employed for the selection of transformants. The introduction of *ptrA* or *hph* into each gene locus was confirmed by PCR using the primer pairs mnn5-F/ptrA-R and ptrA-F/mnn5-R, mnn5-F/hph-R and hph-F/mnn5-R, respectively (Fig. S1).

### Construction of Δ*mnn2*Δ*mnn5*Δ*mnn9*, Δ*mnn2*Δ*mnn5*Δ*van1*, and Δ*mnn2*Δ*mnn5*Δ*anpA* strains

The *mnn9*, *van1*, or *anpA* were disrupted in *A. fumigatus* Δ*mnn2*Δ*mnn5* (*ptrA*) by *hph* insertion. A gene replacement cassette, including the homology arm at the 5′ end of the *mnn9*, *van1* or *anpA* genes and the homology arm at the 3′ end of the *mnn9*, *van1*, or *anpA* genes, was generated by recombinant PCR using *A. fumigatus* A1151 genomic DNA as the template and the primer pairs xxxx-1/xxxx-2, and xxxx-3/xxxx-4, respectively (where “xxxx” indicates *mnn9*, *van1*, or *anpA*) (Table S2). The *hph* marker was amplified by recombinant PCR using pHSG396-hph (Kadooka et al. 2022a) as the template and the primer pairs pHSG396-F/pHSG396-R. The resulting DNA fragment, amplified with primers xxxx-1 and xxxx-4, was used to transform *A. fumigatus* A1160, generating Δ*mnn2*Δ*mnn5*Δ*mnn9*, Δ*mnn2*Δ*mnn5*Δ*van1*, or Δ*mnn2*Δ*mnn5*Δ*anpA* strains, respectively. MM agar plates supplemented with 200 µl/ml hygromycin B were employed for the selection of transformants. The introduction of *hph* into each gene locus was confirmed by PCR using the primer pair xxxx-F/hph-R and hph-F/xxxx-R (Fig. S1).

### Construction of ΔScmnn2ΔScmnn5 in S. cerevisiae

The *Scmnn2* gene was disrupted in *S. cerevisiae* Δ*Scmnn5* by *HIS3* insertion. A gene replacement cassette, consisting of 50 bp of the homology arm at the 5′ end of the *mnn2* gene, *HIS3*, and 50 bp of the homology arm at the 3′ end of the *mnn2* gene, was amplified by PCR using pRS313 plasmid DNA as the template and the primer pairs ScMNN2-HIS3-F/ScMNN2-HIS3-R (Table S2). SC agar plates without uracil and uridine were employed for the selection of transformants. The introduction of *HIS3* into each gene locus was confirmed by PCR using the primer pairs ScMNN2-F/HIS-R and HIS-F/ScMNN2-R.

### Construction of YEp352-Mnn2 and YEp352-Mnn5 plasmids

The expression plasmid YEp352-GAP-II was employed as the vector for expressing Mnn2 and Mnn5 in *S. cerevisiae* strains. It’s worth noting that for Mnn2, the transmembrane region of *Sc*Mnn2p (amino acids 1 to 26) was fused with Mnn2 (amino acids 31 to 493), and for Mnn5, the transmembrane region of *Sc*Mnn5p (amino acids 1 to 35) was fused with Mnn5 (amino acids 30 to 492).

For the construction of YEp352-Mnn2, the DNA fragment encompassing the transmembrane region of *Sc*Mnn2p was PCR-amplified using genomic DNA extracted from *S. cerevisiae* BY4741 strain as the template, with the primer pair ScMnn2p-TM(1-26)-F/ScMnn2p-TM(1-26)-R. Subsequently, the Mnn2 fragment was amplified using genomic DNA from the *A. fumigatus* A1151 strain as the template, with the primer pair YEp352-GAPII-Mnn2-F/YEp352-GAPII-Mnn2-R. Additionally, the 3xFLAG fragment was amplified using the pDC1-3xFLAG plasmid (Komachi et al. 2013) as a template and the primer pair 3xFALG-F/3xFALG-R. These three amplified fragments were then seamlessly integrated into the *Sac* I site of YEP352-GAPII (Oka et al. 2007) using the In-Fusion HD Cloning Kit (Takara). Similarly, for the generation of YEp352-Mnn5, the DNA fragment covering the transmembrane region of *Sc*Mnn5p was PCR-amplified using genomic DNA from *S. cerevisiae* BY4741 strain as the template, with the primer pair ScMnn5p-TM(1-35)-F/ScMnn5p-TM(1-35)-R. Next, the Mnn5 fragment was amplified using genomic DNA from the *A. fumigatus* A1151 strain as the template, with the primer pair YEp352-GAPII-Mnn5-F/YEp352-GAPII-Mnn5-R. The two amplified fragments, together with the 3xFLAG fragment, were cloned into the *SacI* site of YEp352-GAP-II using the In-Fusion HD Cloning Kit. The transformation of budding yeast was performed using the lithium acetate method.

### Preparation of soluble cell wall fraction and ^1^H-NMR analysis

Mannoproteins and galactomannan from *A. fumigatus* were prepared following a previously established method (Chihara et al. 2020). ^1^H-NMR analysis was performed following a previously established method (Chihara et al. 2020). The cell wall extract underwent fractionation by acetyl trimethyl ammonium bromide (CTAB) using a known procedure. The CTAB fraction was precipitated at pH 9.0 with NaOH in the presence of borate, resulting in the recovery of total mannoproteins and FTGM. To remove O-linked glycans, a β-elimination reaction was executed by subjecting the fractionated mannoproteins and FTGM to reducing alkali conditions (500 mM NaBH4/100 mM NaOH, 10 ml, at 25°C for 24 h). Following neutralization with a 50% acetic acid solution, the samples underwent overnight dialysis against distilled water. The purified samples were then lyophilized, resuspended in distilled water, and clarified using 0.45-µm pore filters. For the removal of galactofuran side chains, FTGMs underwent treatment with 100 mM hydrochloric acid at 100°C for 60 min. Subsequently, the samples were neutralized with 10 M NaOH and subjected to overnight dialysis against dH_2_O. In preparation for Nuclear Magnetic Resonance spectroscopy, samples for NMR were exchanged twice in D_2_O with intervening lyophilization. They were then dissolved in D_2_O (99.97% atom 2H).

### Construction of pET15-SmaI plasmid

The adapted plasmid fragments were amplified via PCR, employing pET15-KAI (Katafuchi et al. 2017) as a template with the primer pairs pET15-SmaI-F and pET15-SmaI-R. Subsequently, the amplified DNA fragments were self-ligated using the In-Fusion HD cloning kit (TAKARA, Japan) to generate pET15-SmaI.

### Construction of pET15-Mnn2, pET15-Mnn5, pET15-Mnn9, and pET15-Van1 plasmids

As the *mnn2* and *mnn5* genes lack introns, PCR amplification was conducted using genomic DNA from the *A. fumigatus* A1151 strain as a template, with primer pairs pET15-Mnn2-F, pET15-Mnn2-R, pET15-Mnn5-F, pET15-Mnn5-R, respectively. For the *mnn9* genes, PCR amplification utilized total cDNA prepared from *A. fumigatus* A1151 with the primer pair pET15-Mnn9-F, pET15-Mnn9-R. The resulting DNA fragments were inserted into the *Nde* I and *Not* I sites of pET15-KAI (Katafuchi et al. 2017) using the In-Fusion HD cloning kit, yielding pET15-Mnn2, pET15-Mnn5, and pET15-Mnn9. The *Escherichia coli* codon-optimized *van1* gene fragments from *A. fumigatus* A1163 were synthesized by gBlocks (Integrated DNA Technologies, USA) and cloned into the *Sma* I site of pET15-SmaI using the In-Fusion HD cloning kit, resulting in pET15-Van1. Construction of the plasmid for expressing the C-terminal truncated form (Δ387-499) of *van1* involved PCR amplification of the *van1* (Δ387-499) gene using pET15-Van1 as a template with the primer pair pET15-SmaI-Van1(Δ387-499)-F and pET15-SmaI-Van1(Δ387-499)-R. The amplified DNA fragments were then cloned into the *Sma* I site of pET15-SmaI using the In-Fusion HD cloning kit, yielding pET15-Van1(Δ387-499). Subsequently, the plasmids were transformed into SHuffle T7 Express (New England Biolabs, USA) containing the plasmid pRARE extracted from Rosetta-gami B(DE3) (MilliporeSigma, Burlington, MA).

### Construction of pET15-AfMnn9-AfVan1 (no-tag) plasmid

The *mnn9* or *van1* genes were amplified by PCR using pET15-Mnn9 and pET15-Van1(Δ387-499) as templates, with primer pairs pET15-SmaI-Mnn9-Van1(Δ387-499)(no-tag)-F1, pET15-SmaI-Mnn9-Van1(Δ387-499)(no-tag)-R1, and pET15-SmaI-Mnn9-Van1(Δ387-499)(no-tag)-F2, pET15-SmaI-Mnn9-Van1(Δ387-499)(no-tag)-R2, respectively. The resulting DNA fragments were then inserted into the *Sma* I site of pET15-SmaI using the In-Fusion HD cloning kit, yielding pET15-SmaI-Mnn9-Van1(Δ387-499)(no-tag).

### Protein purification and quantification

Bacterial expression and purification of 6xHis tagged proteins were conducted following a previously described method (Katafuchi et al. 2017).

### Enzyme assays

The synthetic acceptor substrate, *p*-nitrophenyl α-D-mannopyranoside (α-Man-pNP) was procured from Tokyo Chemical Industry Co., Ltd. (Japan). Standard assays were conducted with α-Man-pNP (1.5 mM) as the acceptor, GDP-Man (5 mM) as the donor, and each purified protein (0.2 µg/µL) in a total reaction volume of 40 µL. The reaction mixture was incubated at 30°C for 16 h and terminated by heating at 99°C for 5 min. The reaction mixture was subjected to analysis by HPLC using a Shodex Asahipak NH2P-50 4E amino column (250 × 4.6 mm; Showa Denko, Japan), as previously detailed (Katafuchi et al. 2017). Alpha-(1→2)-mannosidase was obtained by expressing the AfmsdS/AfmsdC gene in *E. coli* (Li et al. 2008).

For Mnn9 and Van1 products, the supernatant underwent analysis by reverse phase-HPLC using an InertSustain C18 column (250 × 4.6 mm; GL Science, Japan), as outlined previously (Kadooka et al. 2022b). Para-nitrophenol derivatives were detected using UV_300_ absorbance. Jack bean mannosidase and α-(1→2, 3) Mannosidase were procured from New England Biolabs (USA).

## Acknowledgments

Strains were obtained from the Fungal Genetics Stock Center (Kansas City, MO). This work was supported in part by Grant-in-Aid for Scientific Research (C) from the Japan Society for the Promotion of Science (JSPS KAKENHI) (18K05418 to T.O., 21K05373 to T.O., and 22K06600 to Y.T.), Grant-in-Aid for Early-Career Scientists from Japan Society for the Promotion of Science (JSPS KAKENHI) (22K14817 to C.K., and 20K15997 to Y.T.).

We declare that the research was conducted in the absence of any commercial or financial relationships that could be construed as a potential conflict of interest.

C.K., R.K., and T.O. performed the experiments. Y.T. performed NMR analysis. C.K., D.H., Y.T., and T.O. analyzed and interpreted the data. T.O. conceived and designed the research project. C.K., and T.O. wrote the manuscript. All authors contributed to the article and approved the submitted version.

## Figure Legends

**Fig. S1.**
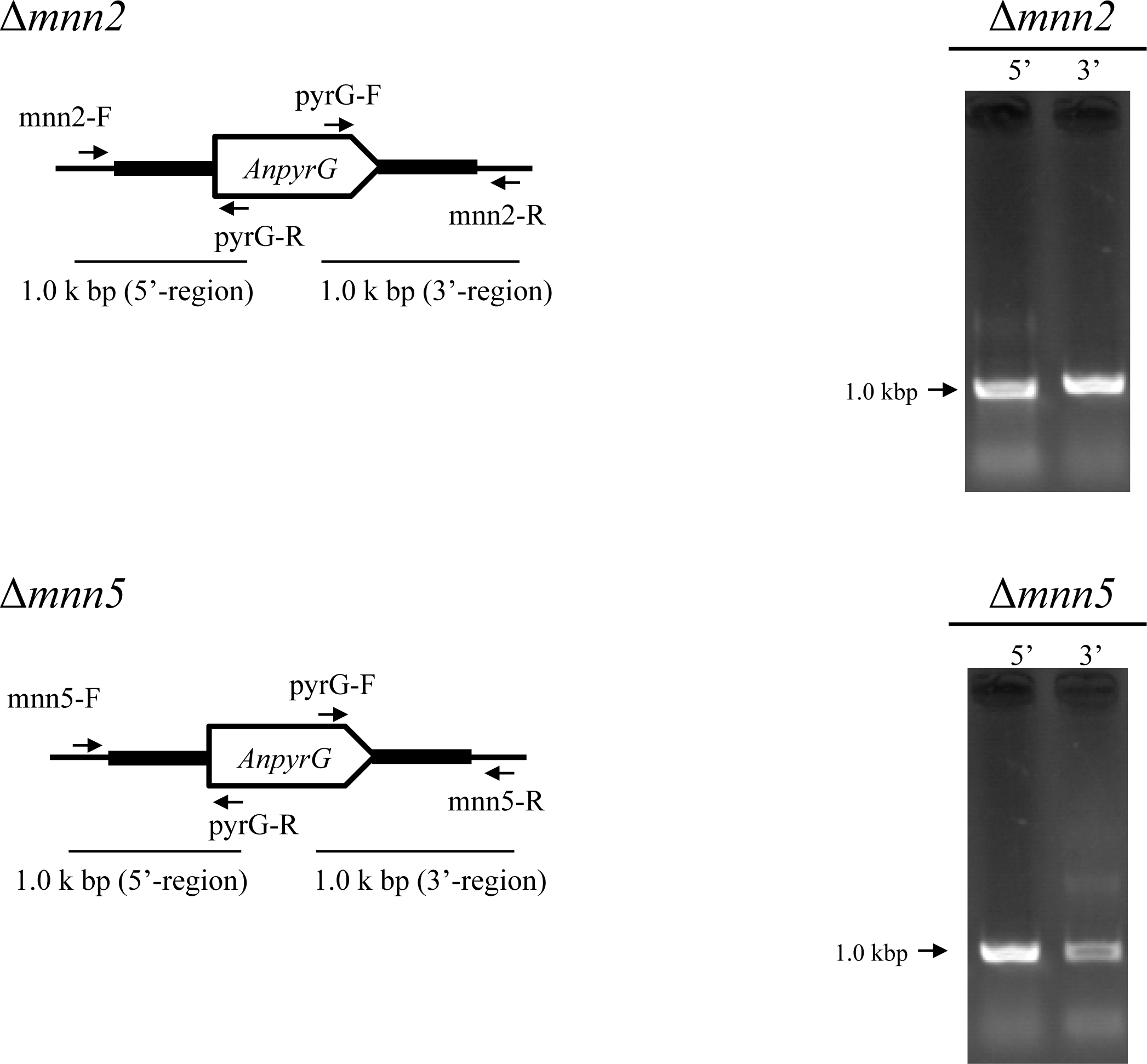
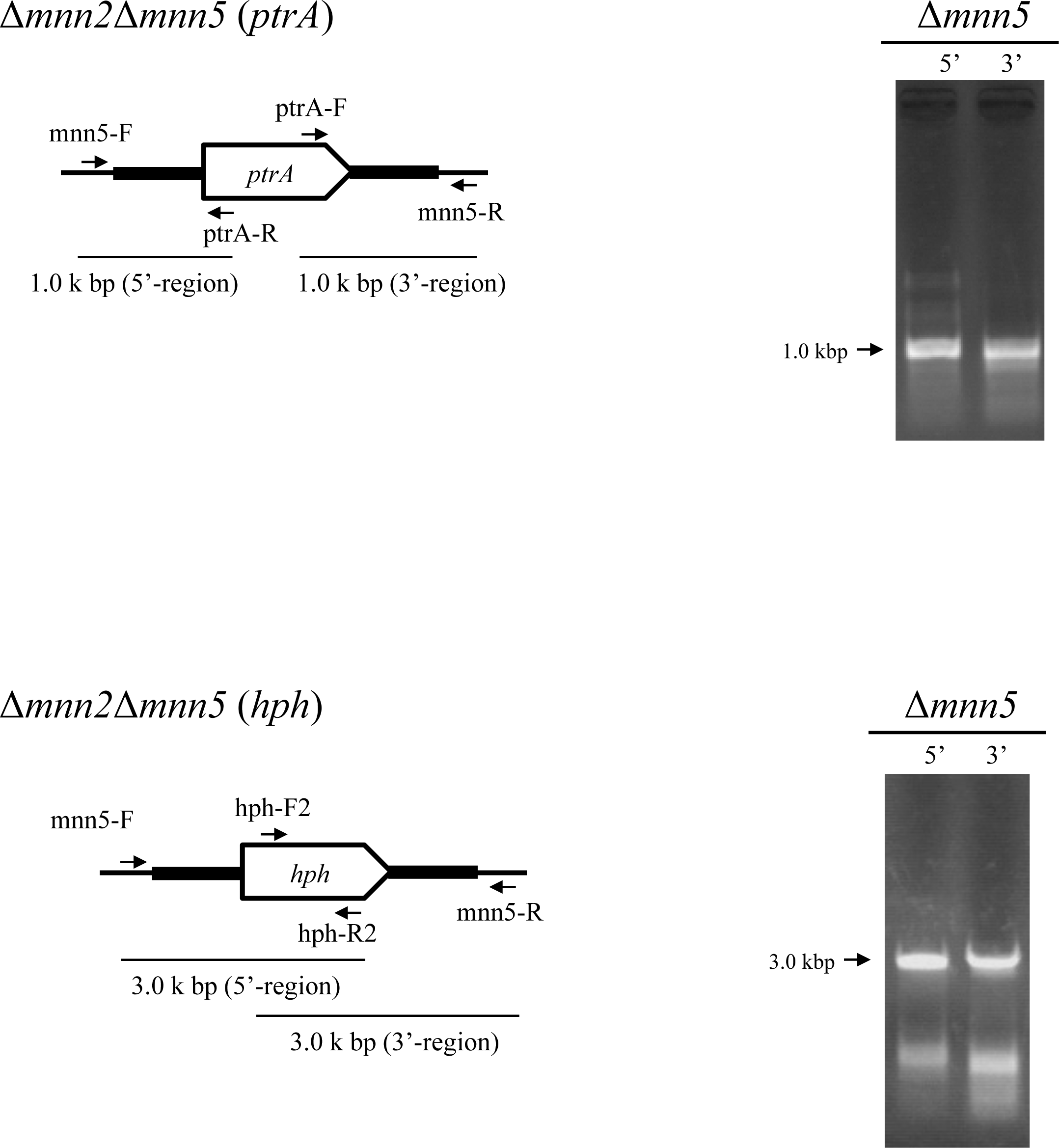
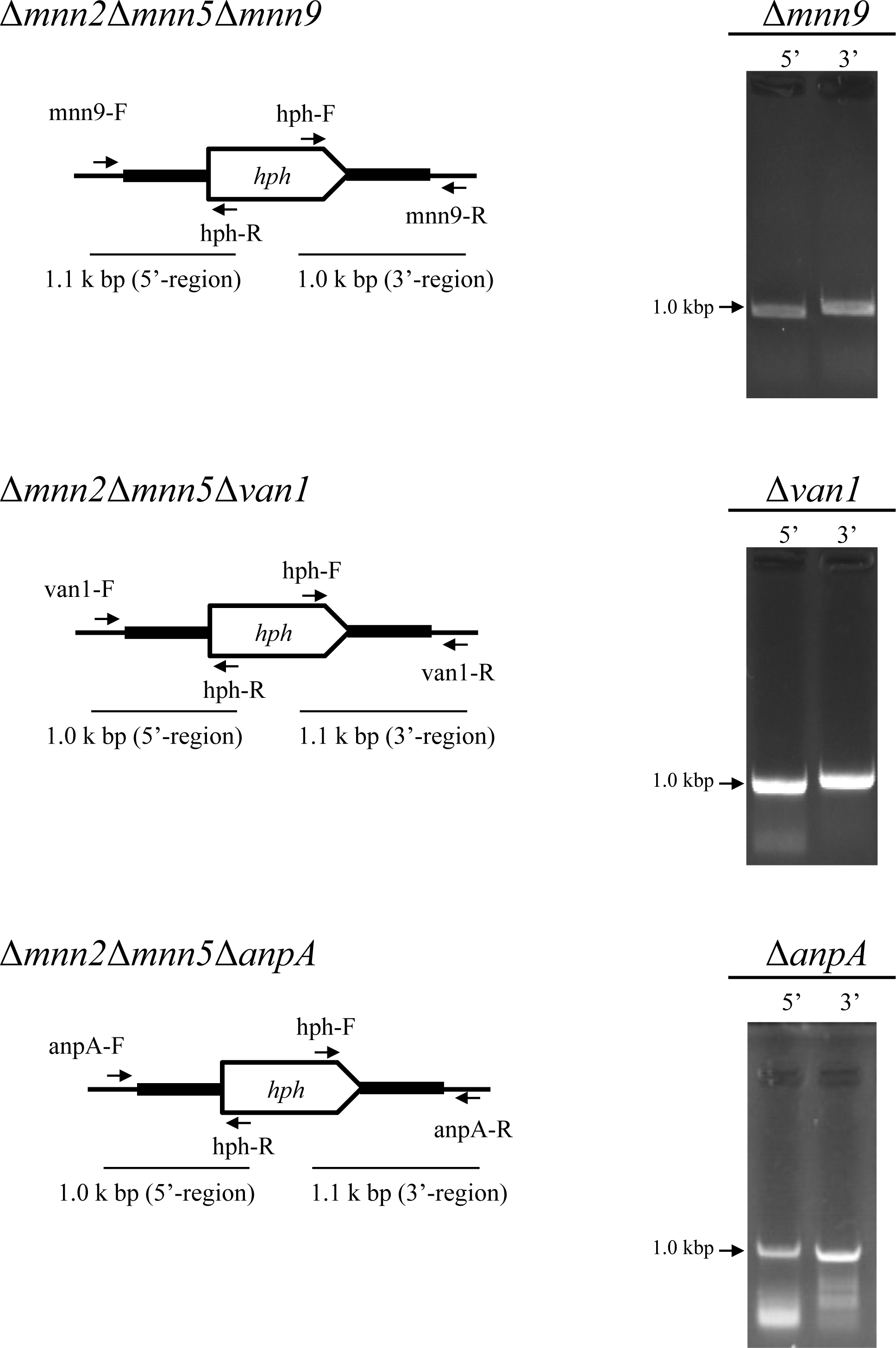
Construction of *A. fumigatus* strains used in this study. Disruption of *mnn2*, *mnn5, mnn9, van1*, double disruption of *mnn5* in Δ*mnn2*, triple disruption of *mnn9*, *van1*, and *anpA* in Δ*mnn2*Δ*mnn5* (*ptrA*), respectively. Results of electrophoretic analyses of the PCR products are shown in the panels to the below.

**Table S1.**
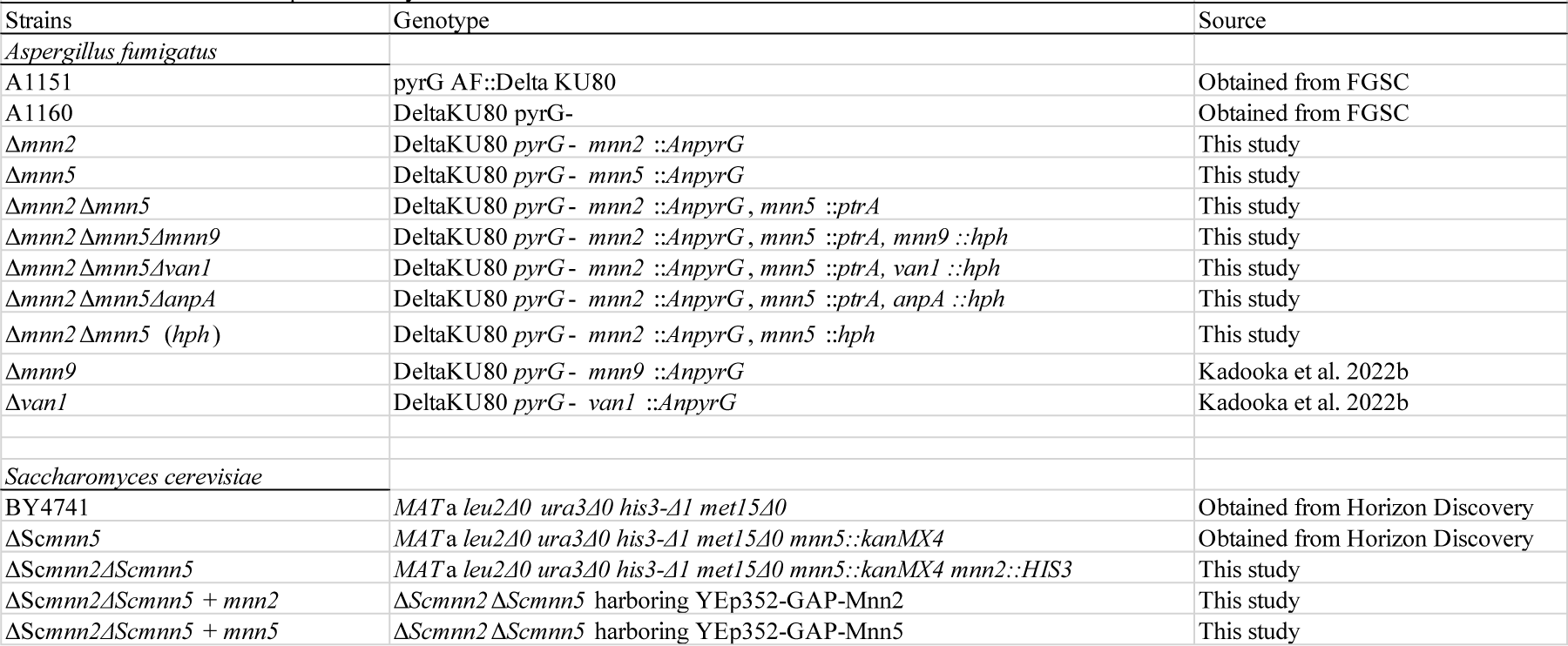
Strains used in the present study.

**Table S2.**
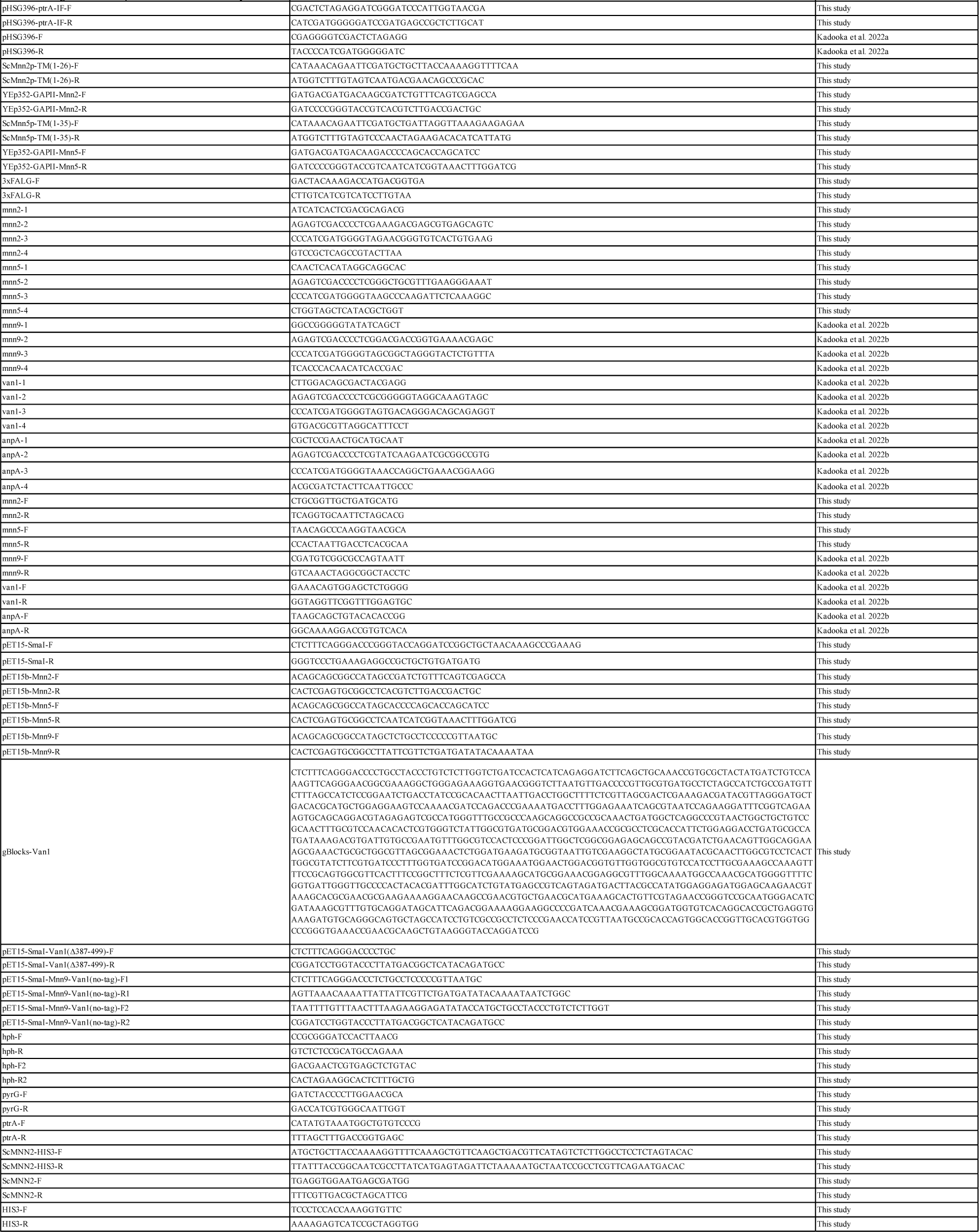
Oligonucleotide primers used in this study.

